# Crowding-induced collapse and adsorption of polymers with nonuniform bending stiffness

**DOI:** 10.1101/2025.09.04.674235

**Authors:** Gregory R. Cantrall, Gaurav Chauhan, Steven M. Abel

**Author notes:** Department of Chemical Engineering, Indian Institute of Technology, Indore (Gaurav Chauhan).

## Abstract

Macromolecular crowding can significantly impact the behavior of biopolymers, with crowding-induced depletion interactions influencing both the conformations and surface adsorption of individual polymers. Although previous studies have explored the influence of homogeneous polymer stiffness in crowded conditions, biomolecules such as DNA can exhibit sequence-dependent stiffness, and DNA origami nanoparticles can be designed with alternating stiff and flexible domains. In this work, we use Langevin dynamics simulations to characterize how nonuniform bending stiffness modulates the conformations and adsorption of polymers in crowded environments. By systematically varying the relative length and arrangement of flexible and semiflexible domains along a linear chain, we show that increasing osmotic pressure leads to a pattern-dependent collapse of the polymer, as revealed by a decrease in the radius of gyration. In general, large flexible regions promote polymer collapse, although flexible domains separating extended semi-flexible regions can facilitate their contact, leading to stable folded conformations. When a surface is present, large semiflexible domains promote adsorption, and the pattern of stiffness can be used to control the adsorption threshold. Our findings provide insight into the impact of spatially varying stiffness on the behavior of polymers in crowded environments, highlighting mechanisms relevant to biopolymers and deformable nanoparticles in both cellular and cell-free contexts.

## 1 Introduction

Macromolecular crowding is common in cells, with up to 40% of the volume of the cytosol occupied by macromolecules. ^1,2^ One important consequence of crowding is the emergence of attractive depletion interactions,^3,4^ which can affect cellular organization ^2,5^ and have been implicated in genome organization, ^6–9^ gene regulation, ^10^ and intracellular phase separation. ^11,12^ Macromolecular crowding can also be utilized in cell-free synthetic systems to manipulate the organization of biomolecules such as DNA plasmids. ^13,14^

Understanding the behavior of polymers in crowded environments is of particular interest because biopolymers such as DNA, actin, and microtubules are susceptible to effects of depletion interactions, which can cause both intra- and inter-polymer attraction. Experimental studies have shown that molecular crowding can cause compaction of DNA, ^15,16^ the condensation of isolated actin filaments, ^17^ and aggregation of DNA plasmids. ^18^ Computer simulations have revealed how crowding leads to the compaction of linear polymers, ^19^ the collapse of a model chromosome, ^20^ and attraction between ring polymers. ^21^

Macromolecular crowding can also induce attractive depletion interactions between biopolymers and surfaces, thus promoting their adsorption. ^22^ Crowding has been shown to drive DNA plasmids to localize near vesicle surfaces, ^14,23,24^ and the level of crowding can be tuned to either partially or completely adsorb actin filaments. ^25^ Simulations of flexible ring polymers have demonstrated that polymer-polymer attraction is markedly weaker than polymer-surface attraction, with effective polymer-surface interactions emerging at lower levels of crowding. ^21^ Crowding can also cause confined model polymers to localize near confining surfaces. ^26–28^

The adsorption of polymers onto surfaces due to short-ranged attractive interactions has been of long-standing interest in polymer physics, with significant efforts devoted to understanding the behavior of both flexible and semiflexible polymers. ^29,30^ Semiflexible polymers, which are characterized by a nonzero bending stiffness, are of particular interest in biology, as biopolymers such as DNA, actin filaments, and microtubules are characterized by distinct stiffnesses. The bending stiffness of a polymer is an important factor in controlling its adsorption because adsorption leads to a loss of conformational entropy, and stiffer polymers lose less conformational entropy upon adsorption. Thus, with other factors held constant, increasing the stiffness of a polymer increases its propensity to adsorb. ^31–38^

While significant progress has been made in understanding flexible and semiflexible polymers with uniform mechanical properties, much less is known about polymers with nonuniform bending stiffness. DNA, for example, exhibits sequence-dependent bending stiffness, underscoring the relevance of local variation. ^39–42^ High-throughput studies of long DNA sequences have shown that the persistence length of double-stranded DNA increases with increasing fraction of GC content,^41^ which is consistent with detailed analysis of specific short DNA sequences. ^40^ Additionally, the combination of flexible and semiflexible domains can influence the self-assembly of synthetic copolymers. ^43^

Understanding the effects of nonuniform mechanical properties is also relevant in DNA nanotechnology, ^44^ where deformable DNA origami structures with regions of differing stiffness have generated interest. ^45–48^ A common motif is a hinge-like DNA origami structure, where two relatively rigid arms are connected by a short region whose stiffness can be tuned. ^45,46^ Simulations demonstrate that when interacting with membrane surfaces through short-ranged attraction, the stiffness of the hinge region impacts the free energy of binding to the membrane and the self-assembly of particles on the surface. ^49–51^ Other mechanically-compliant DNA origami designs have incorporated multiple hinges with controlled location, stiffness, and angle, ^52^ introduced engineered defects to locally control flexibility, ^53^ and incorporated single-stranded DNA to program bending and flexibility. ^54^ Structural dynamics of a DNA device have also been designed to switch between conformational states in a manner that allows measurement of depletion forces. ^55^

Despite the relevance of nonuniform bending stiffness in both biopolymers and deformable nanostructures, little is known about how such systems are impacted by attractive depletion interactions. In this study, we employ Langevin dynamics simulations to explore polymers with alternating flexible and semiflexible segments. We systematically vary the relative lengths of the regions, their spatial arrangement along the polymer, and the osmotic pressure of the environment.

We first characterize the conformations and radius of gyration of isolated polymers, and then examine their adsorption onto surfaces. Together, our results reveal how different patterns of bending stiffness modulate the compaction and adsorption of polymers in crowded environments, providing insights relevant to both biopolymers and deformable nanostructures in cellular and cell-free systems.

## 2 Methods

We used Langevin dynamics simulations to investigate the behavior of isolated polymers with nonuniform bending stiffness in crowded environments. We considered polymers of fixed length consisting of both flexible and relatively stiff segments. We varied the relative length and spatial distribution of each type of segment. The impact of macromolecular crowding was accounted for using implicit depletion interaction potentials, and we varied the osmotic pressure to investigate the impact of varying levels of crowding. Simulations were conducted both with and without the presence of a surface.

We utilized a bead-spring polymer model consisting of *N* = 50 beads. Adjacent beads were connected by a finitely extensible nonlinear elastic (FENE) bond potential,

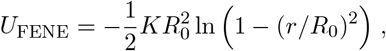

where *R*_0_ = 1.5*σ* represents the maximum distance between two adjacent beads, *r* denotes the center-to-center distance between the beads, and *K* = 15*ϵ/σ*^2^ is the spring constant. *σ* and *ϵ* set the distance and energy units, respectively. The bending potential for each set of three consecutive beads was

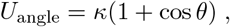

where *κ* is the bending stiffness and *θ* is the angle formed by the beads. Each polymer consisted of flexible and semiflexible segments characterized by a different bending stiffness: We set *κ* = 0 for flexible segments and *κ* = 50*ϵ* for semiflexible segments. Given that the polymer consisted of 50 beads, a total of 48 angles contributed to the bending energy. We let *η* denote the fraction of flexible segments. The number of flexible segments varied from 0 (*η* = 0) to 48 (*η* = 1), corresponding to polymers that ranged from purely semiflexible to purely flexible.

The beads of the polymer also interacted through depletion interactions and purely repulsive short-range interactions. The implicit depletion potential between each pair of beads was^4,56,57^

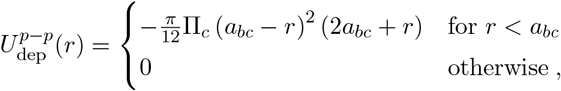

where Π_*c*_ is the osmotic pressure, *a*_*bc*_ = *a*_*c*_ + *a*_*b*_, *a*_*b*_ is the diameter of the polymer beads, and *a*_*c*_ is the diameter of the crowders. Each pair of beads also interacted through the purely repulsive Weeks-Chandler-Andersen potential, ^58^

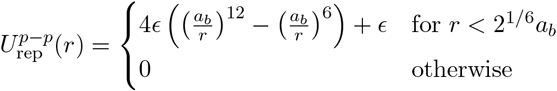

When a surface was present, the interaction of each polymer bead with the surface was given by an attractive depletion interaction and a short-ranged repulsive interaction, ^4^

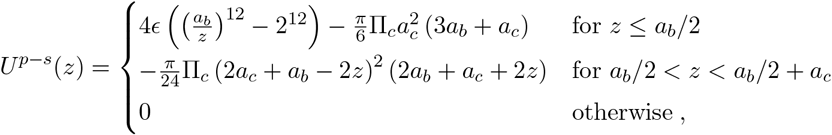

where *z* denotes the distance between the bead and the surface. In this work, both the diameter of the polymer beads and the diameter of the crowders were fixed at *a*_*b*_ = *a*_*c*_ = *σ*. To simulate environments with different levels of crowding, we systematically varied the osmotic pressure (Π_*c*_).

We considered four patterns for the flexible domains. The *diblock* pattern consisted of a single flexible region adjoined to a single semiflexible region (Figure 1A). The *centered* arrangement consisted of a continuous flexible region in the center of the polymer, with equally-sized semiflexible domains on each side (Figure 1B). The *uniform* pattern evenly distributed flexible regions throughout the polymer, creating multiple flexible and semiflexible domains (Figure 1C). For a given value of *η*, the flexible regions were distributed to create multiple semiflexible regions of approximately the same length. In the *random* arrangement, a specified number of angles along the polymer were chosen at random to be flexible, while the others had nonzero bending stiffness (Figure 1D). Polymers with different arrangements exhibit different behaviors, so we analyzed four independent random arrangements for each number of flexible segments.

**Figure 1:**
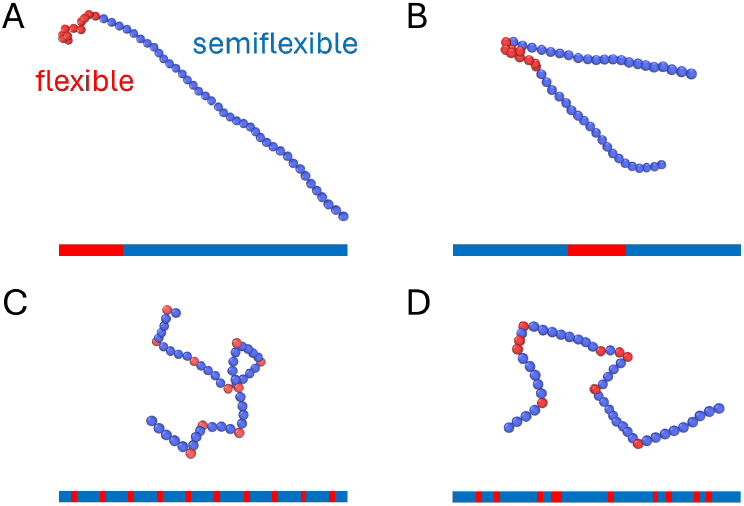
Snapshots of simulated polymers and schematics depicting different patterns of flexibility along the polymer with *η* = 0.21, where *η* denotes the fraction of flexible segments. Blue indicates a semiflexible domain (*κ >* 0), while red indicates a flexible domain (*κ* = 0). (A) Diblock. (B) Centered. (C) Uniform. (D) Random.

Polymers were simulated in a cubic simulation domain with a side length of 100*σ*. For simulations without a surface, periodic boundaries were used in all directions. For simulations with a surface, surfaces were introduced at the edges of the *z*-dimension, with periodic boundary conditions in the *x*- and *y*-dimensions.

Simulations were conducted in LAMMPS using the velocity-Verlet algorithm to integrate the Langevin equations. The timestep was 0.0005*τ*, where *τ* is the simulation time unit. Each simulation started with an equilibration period of 5 × 10^4^*τ*, followed by a production run of 1.75 × 10^5^*τ*. Data was collected at a sampling interval of 50*τ*. Averaged quantities represent a time average over a production run at given conditions.

In the simulations, we independently varied the number of flexible angles, their organization, and Π_*c*_. For each pattern, we considered 0, 5, 10, 20, 30, 40, and 48 flexible segments, which we report in terms of *η*, the fraction of the total number of angles (*η* = 0, 0.10, 0.21, 0.42, 0.63, 0.83, and 1). We considered osmotic pressures of Π_*c*_ = 0, 0.1, 0.2, 0.3, 0.4, 0.5, 0.6, 0.8, and 0.9 *ϵ/σ*^3^, where Π_*c*_ = 0 corresponds to an absence of macromolecular crowding.

## 3 Results and discussion

### 3.1 Flexible regions promote the collapse of polymers in crowded conditions in a pattern-dependent manner

We first consider isolated polymers without a surface and quantify their radius of gyration (*R*_*g*_), defined by

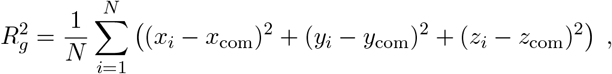

where *N* is the total number of beads, *x*_*i*_, *y*_*i*_, and *z*_*i*_ are the coordinates of bead *i*, and *x*_com_, *y*_com_, and *z*_com_ are the coordinates of the center of mass of the polymer. For a given polymer, we normalize the average radius of gyration at osmotic pressure Π_*c*_ by the average value in the absence of crowding, ⟨*R*_*g*_(Π_*c*_)⟩*/*⟨*R*_*g*_(0)⟩. This metric allows us to assess how crowding influences polymer compaction and later provides insight into adsorption behavior.

We first consider the diblock pattern, in which a continuous flexible domain is connected to a continuous semiflexible domain. When the diblock polymer has a short flexible domain (*η* = 0.10, 0.21), it behaves similarly to the semiflexible polymer (*η* = 0) and exhibits almost no change in ⟨*R*_*g*_⟩ as Π_*c*_ is varied (Figure 2A). In contrast, when the diblock polymer has a large flexible domain (*η* = 0.83), it behaves similarly to the purely flexible polymer (*η* = 1) and exhibits a pronounced reduction in ⟨*R*_*g*_⟩, which decreases by ≈ 80% over the range of Π_*c*_ considered. The conformational changes responsible for these differences are evident in representative equilibrium snapshots (Figure 2D). The semiflexible segment remains extended for diblock polymers even at high crowding levels, while the flexible region collapses into a compact conformation. Thus, when *η* is small, the impact of crowding on ⟨*R*_*g*_⟩ is small because it is dominated by the semiflexible domain. These results highlight that localized flexibility facilitates collapse while semiflexible domains resist compaction, maintaining an extended structure despite the presence of depletion interactions.

**Figure 2:**
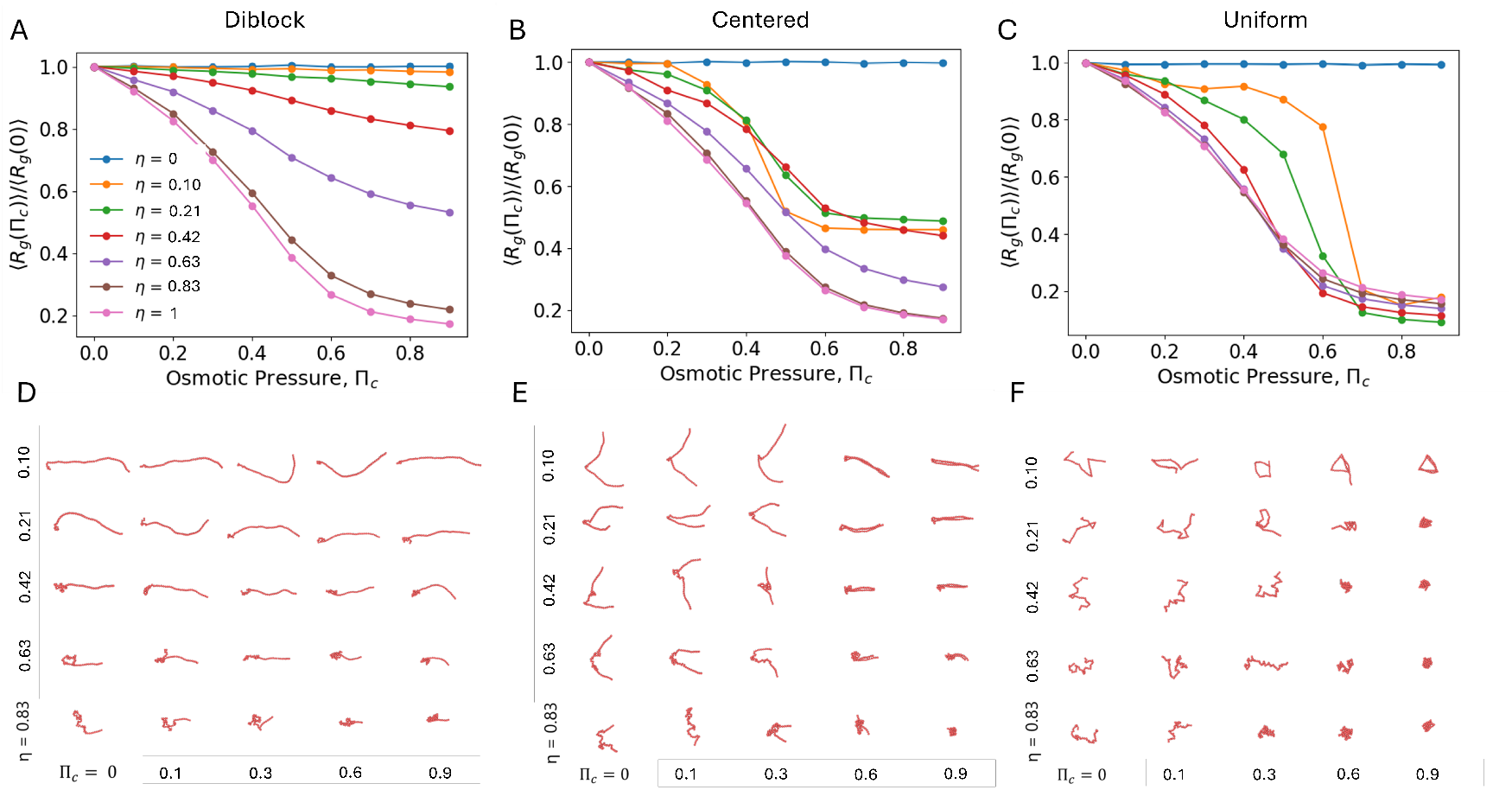
(A - C) Mean radius of gyration in bulk normalized by its mean value at Π_*c*_ = 0 for the diblock (A), centered (B), and uniform patterns (C). The normalized radius of gyration is shown for the fraction of flexible segments ranging from *η* = 0 (semiflexible polymer) to *η* = 1 (flexible polymer). (D - F) Snapshots of diblock (D), centered (E), and uniform (F) patterns. Snapshots are shown on a consistent scale to facilitate direct comparison.

The centered pattern is characterized by two semiflexible domains connected by a single flexible domain. In contrast with the diblock case, even short flexible regions result in marked compaction of the polymer as Π_*c*_ increases (Figure 2B). The normalized radius of gyration plateaus near the same value for flexible domains of short and intermediate length (*η* = 0.10, 0.21, 0.42). Equilibrium snapshots show that the flexible region acts as a hinge that allows the two semiflexible regions to come into contact in a hairpin-like configuration (Figure 2E). Depletion interactions stabilize the contact of the semiflexible domains and hence stabilize the folded configuration starting at intermediate values of Π_*c*_. Polymers with larger flexible domains (*η* ≥ 0.63) exhibit a more pronounced decrease in ⟨*R*_*g*_⟩ with increasing Π_*c*_. In this regime, the decrease in ⟨*R*_*g*_⟩ is more strongly impacted by the globular collapse of the large flexible domain, although snapshots reveal that the semiflexible domains still maintain stable contact at *η* = 0.63.

The uniform pattern has discrete flexible domains distributed along the polymer. These domains can act as hinges in a manner similar to the centered case, allowing semiflexible domains to come close enough to experience attractive depletion interactions. There is a pronounced decrease in ⟨*R*_*g*_⟩ for all cases with *η >* 0, and compared to the diblock and centered patterns, the uniform case leads to the most pronounced decrease of ⟨*R*_*g*_⟩ for polymers with a small fraction of flexible segments (Figure 2C). The cases with small *η* “fold” into structures that allow semiflexible domains to contact one another, such as the triangular configuration observed for *η* = 0.10 (Figure 2F). At intermediate value of Π_*c*_, the contact between segments is transient, but the structures are relatively stable at large values of Π_*c*_. Larger fractions of flexibility promote compact structures at high crowding levels that are similar to those observed for the fully flexible polymer.

To compare the different patterns of flexibility, Figure 3 shows the normalized mean radius of gyration for each pattern at the highest level of crowding, Π_*c*_ = 0.9. The different patterns of flexibility exhibit notable differences as *η* is varied. The diblock pattern exhibits a monotonic decrease in the normalized radius of gyration as *η* increases. In contrast, the centered pattern shows a more pronounced initial decrease, with a sharp reduction at small values of *η*. As *η* increases further, the normalized radius of gyration increases slightly before again decreasing with increasing size of the flexible domain. Polymers with uniform and random patterns behave similarly, showing a marked decrease in the normalized radius of gyration at *η* = 0.10, with the decrease in the uniform case more pronounced. This indicates significant collapse when a small number of flexible regions are distributed throughout the polymer. As *η* increases further, the normalized radius of gyration first decreases before slightly increasing at large *η*.

**Figure 3:**
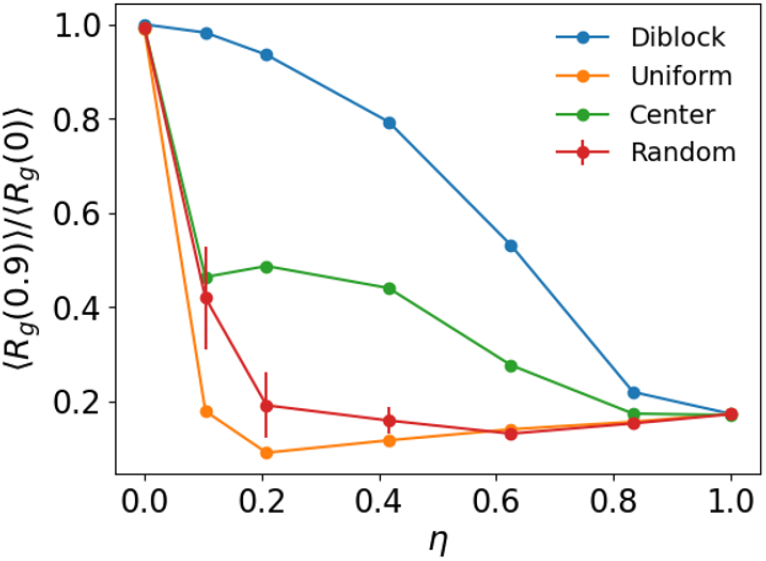
Normalized mean radius of gyration at Π_*c*_ = 0.9 for each pattern of flexibility. Error bars for the random case represent the standard deviation across four independent replicates.

### 3.2 Flexible regions suppress adsorption in diblock polymers

To investigate polymer adsorption, we studied polymers confined between two parallel surfaces at various crowding levels. For the diblock polymer at small values of *η*, there is an increase in the normalized radius of gyration around Π_*c*_ = 0.4 (Figure 4A), which is in contrast with the monotonic behavior observed for polymers without a surface. The increase in ⟨*R*_*g*_⟩ at an intermediate value of Π_*c*_ becomes less pronounced as *η* increases, and ⟨*R*_*g*_⟩ for large values of *η* is similar to the results without a surface.

**Figure 4:**
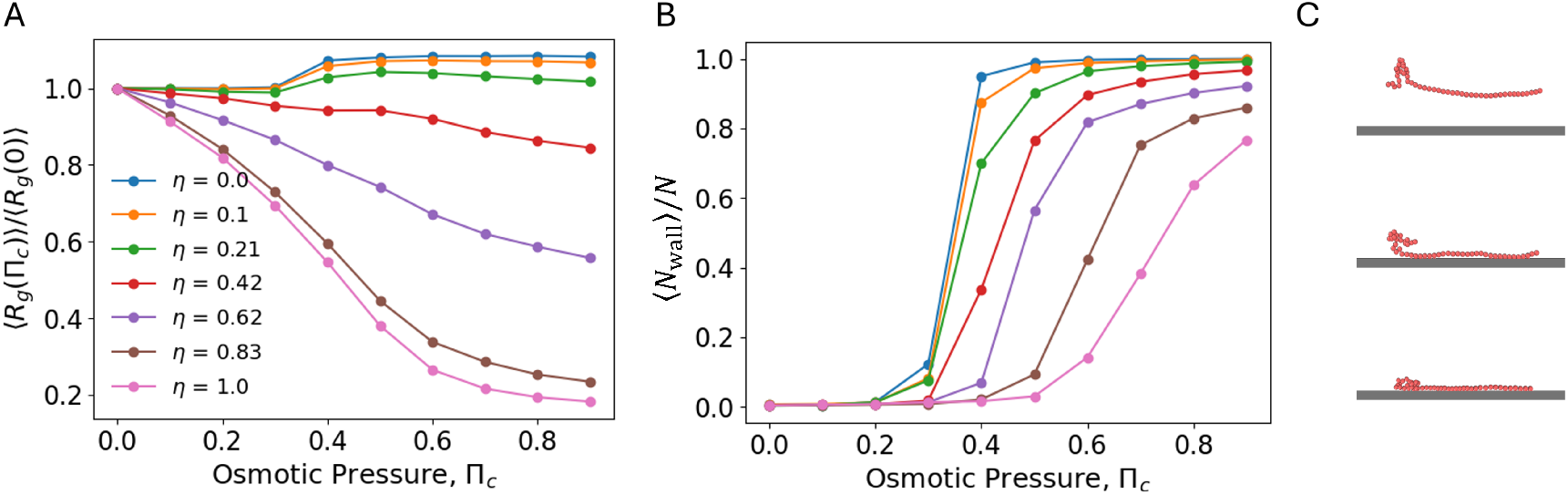
Results for diblock polymers in the presence of a surface. (A) The normalized mean radius of gyration for polymers with varying fractions of flexible segments (*η*). (B) The average fraction of the polymer beads in contact with a surface. (C) Snapshots of a diblock polymer (*η* = 0.21) at Π_*c*_ = 0 (top), 0.4, and 0.9 (bottom).

We further characterize the polymers in terms of the average fraction of beads in contact with a surface, ⟨*N*_wall_⟩*/N*, where *N*_wall_ denotes the number of polymer beads in close apposition to a surface (*z <* 0.65*σ*). The diblock polymers do not adsorb at small values of Π_*c*_, which is reflected by ⟨*N*_wall_⟩*/N* ≈ 0 (Figure 4B). However, for small values of *η*, there is a sharp increase in the fractional contact at Π_*c*_ = 0.4, indicating strong association with a surface for Π_*c*_ ≥ 0.4. The fractional contact plateaus near 1, indicating persistent contact of all polymer beads with the surface. The marked change in fractional contact at Π_*c*_ = 0.4 corresponds to the increase observed in ⟨*R*_*g*_⟩, which is a consequence of the polymer having fewer spatial degrees of freedom, resulting in a larger radius of gyration in a plane parallel to the surface. ^28^ As *η* increases, the transition to larger values of ⟨*N*_wall_⟩*/N* becomes broader and shifts to larger values of Π_*c*_. Taken together, the results in Figures 4A and 4B indicate that flexible domains resist adsorption, which is consistent with previous studies demonstrating that purely semiflexible polymers adsorb more readily than purely flexible polymers.

Figure 4C illustrates the distinct adsorption behaviors of the semiflexible and flexible domains. The semiflexible region drives adsorption at intermediate values of Π_*c*_, making contact with the surface along its length. In contrast, the flexible domain remains farther away from the surface until higher osmotic pressures, where it collapses to maintain extended contact with the surface. This behavior arises because of the entropic cost associated with maintaining surface contact.

Semiflexible domains, which have less conformational entropy than flexible domains, experience a smaller entropic penalty upon adsorption. In contrast, adsorption imposes a larger entropic penalty for flexible domains by reducing the available degrees of freedom and restricting accessible conformations.

### 3.3 The pattern of flexibility impacts polymer adsorption

To further characterize how the pattern of flexibility impacts adsorption, we determined the adsorption threshold, defined as the osmotic pressure required for the average fraction of beads in contact with the wall to reach 0.5. The adsorption threshold increases monotonically with *η* for each pattern of flexibility, but the pattern changes the shape of the curve (Figure 5). The diblock pattern requires the lowest osmotic pressure to adsorb. The centered pattern has the same general shape as the diblock case but with a slightly higher adsorption threshold. Both the uniform and random patterns require higher osmotic pressures to adsorb than the diblock or centered patterns. At high and low values of *η*, the uniform pattern adsorbs more readily than the random pattern, while for intermediate values of *η*, it is more resistant to adsorption. The overall trends are consistent with long continuous semiflexible regions promoting adsorption. For each value of *η*, the diblock pattern has the longest continuous semiflexible region, followed by the centered pattern. The uniform and random cases have substantially shorter continuous semiflexible regions, thus requiring higher osmotic pressures to adsorb.

**Figure 5:**
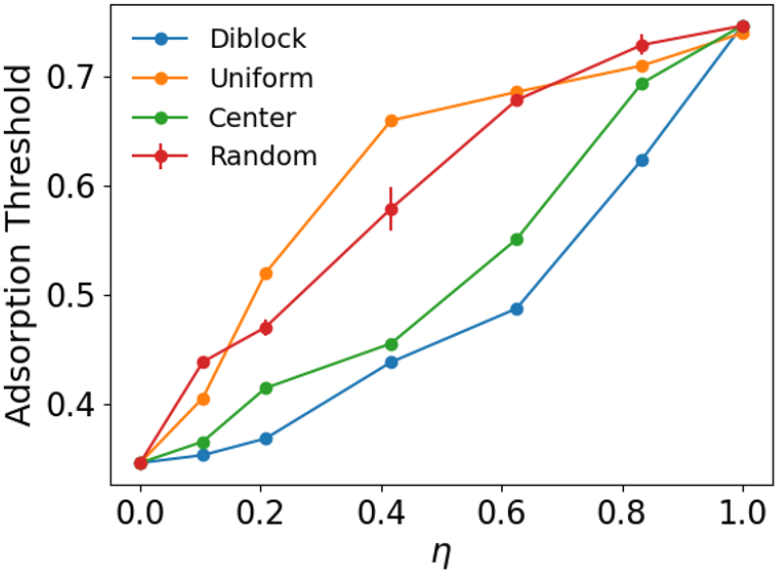
The adsorption threshold, defined as the osmotic pressure required for the average fraction of beads in contact with the wall to reach 0.5. Each pattern of flexibility is shown. Error bars for the random case represent the standard deviation across four independent replicates.

The previous results demonstrate that altering the pattern of flexibility can affect both the collapse of a polymer and its adsorption. In Figure 6, we characterize how the pattern of flexibility influences the adsorption of specific regions of the polymer. For the diblock and centered patterns, the flexible region maintains less contact with the wall than the semiflexible regions (Figures 6A and 6B). In contrast, for the uniform case, each isolated flexible domain does not significantly reduce the contact of a particular region (Figure 6C). Instead, the distributed flexibility leads to lower average adsorption, which is consistent with the adsorption thresholds observed in Figure 5. In all cases, there is a local decrease in contact with the wall at the ends of the polymer, which is consistent with previous studies of partially adsorbed semiflexible polymers. ^25^ This is a consequence of partial desorption at the ends of the polymer being entropically more favorable than near the center.

**Figure 6:**
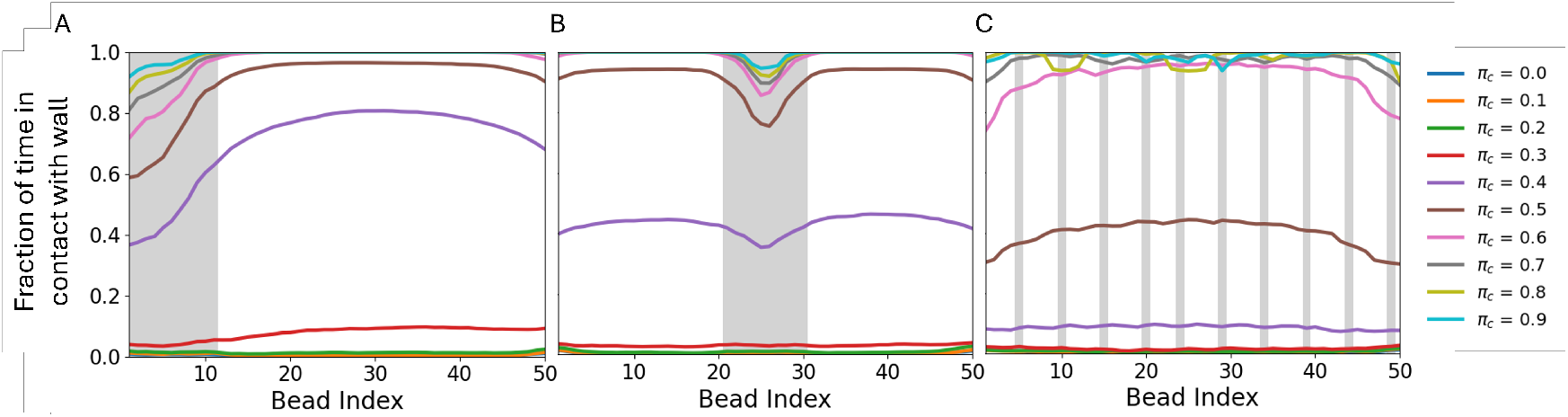
The fraction of time each bead is in contact with a surface. Gray regions indicate flexible domains (*η* = 0.21). (A) Diblock. (B) Centered. (C) Uniform.

Typical conformations adopted by different polymers at large values of Π_*c*_ are shown in Figure 7. Both the diblock and centered patterns adopt simple conformations because they have a single flexible domain. In the diblock polymers, the semiflexible domain remains largely unchanged and extended, while the flexible domain collapses. This can be observed in snapshots and in the average distance map demonstrating close proximity of beads in the flexible region (Figure 7A and 7D). The centered pattern facilitates interactions between the two semiflexible domains, with the flexible domain serving as a hinge that allows the semiflexible domains to come into close contact (Figure 7B and 7E). The anti-diagonal feature in the average distance map indicates strong contact between the two semiflexible domains. In contrast, the uniform pattern allows each of the small flexible domains to act as hinges, resulting in a more complex structure in which different continuous semiflexible domains can come into contact. In the case shown, the polymer collapses into a condensed structure (Figures 7C and 7F), consolidating the stiff domains into a conformation that permits adsorption with surface-bead contact between most polymer beads. The linear features in the average distance map highlight contact between different segments of the polymer.

**Figure 7:**
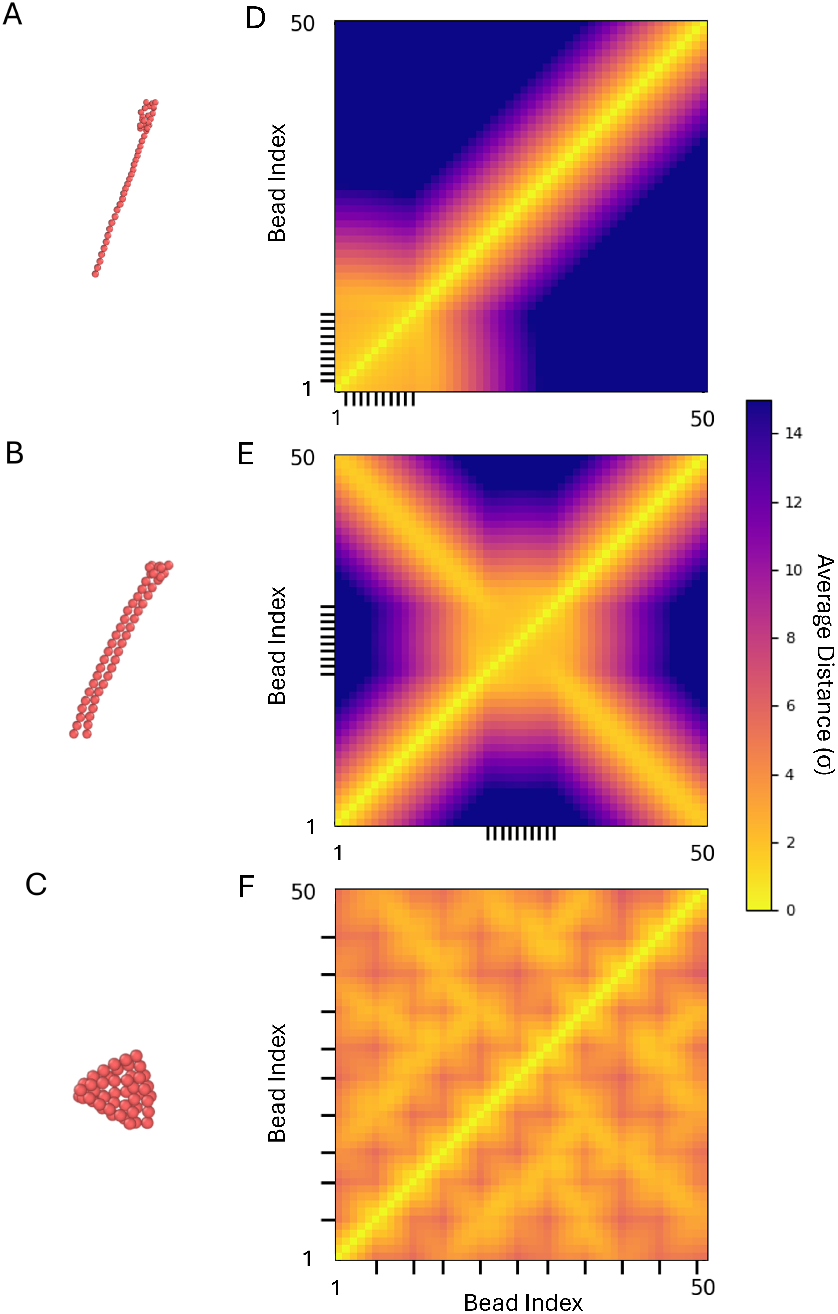
(A-C) Snapshots of polymers with different patterns of flexibility (*η* = 0.21, Π_*c*_ = 0.9). (D-F) Heat maps indicating the average distance between the beads of the polymers. Dashes on the perimeter indicate the location of flexible hinges on the polymer. (A, D) Diblock. (B, E) Centered. (C, F) Uniform.

### 3.4 Variation in the collapse and adsorption of polymers with randomly distributed flexible segments

Randomly distributing flexible domains can lead to variation in behavior, even at fixed *η*, due to differences in their location along the polymer. The variation is most pronounced for the radius of gyration with small numbers of flexible regions (*η* ≤ 0.21), as illustrated for fixed *η* in Figure 8. For larger values of *η*, the differences between difference arrangements of flexibility are less pronounced, and the curves are nearly indistinguishable for *η* ≥ 0.63. Physically, the origin of the differences at small *η* is that the polymers can collapse into different structures that are determined by the lengths of the continuous semiflexible domains between flexible hinges. At larger values of *η*, the impact of the size and location of any specific semiflexible domain becomes less significant, leading to a more homogeneous collapse between the randomly generated patterns. Interestingly, the large variation in ⟨*R*_*g*_⟩ observed at small values of *η* is not reflected in the adsorption behavior of the polymers (Figure 8). While some polymers exhibit minor differences, randomly distributing the flexible regions results in similar average contact, even though the polymers collapse in different ways.

**Figure 8:**
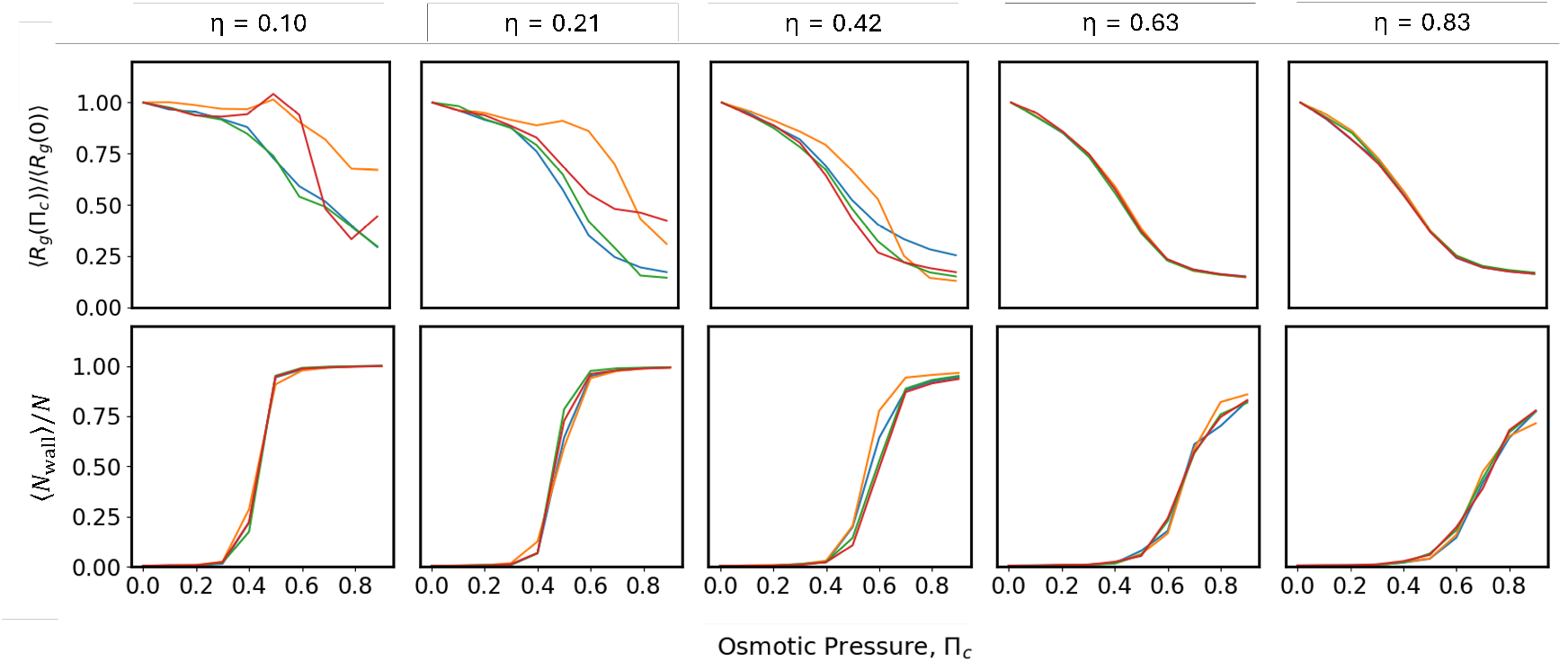
Normalized mean radius of gyration (top) and fractional contact (bottom) for polymers with randomly distributed hinges. Four cases, each with a different random pattern of flexibility, are shown for each value of *η*.

## 4 Conclusions

In this work, we used Langevin dynamics simulations to investigate the collapse and adsorption of polymers with nonuniform bending stiffness in crowded conditions. The relative total length of flexible and semiflexible domains, as well as their spatial pattern along the contour length of the polymer, dictate the extent of polymer compaction and the propensity for surface adsorption with increasing osmotic pressure. While previous studies have explored the collapse and adsorption of flexible and semiflexible polymers, their focus was on polymers with uniform mechanical properties. Our work extends this by introducing nonuniform bending stiffness as a new physical feature. Our findings are consistent with previous work showing that semiflexible polymers, when compared to flexible polymers, resist collapse and have a greater propensity to adsorb. However, our results reveal significant nontrivial effects arising from nonuniform bending stiffness.

Figure 2 reveals key features of the collapse behavior. In general, flexible regions promote collapse. For the diblock pattern, the two regions exhibit distinct behavior; the flexible region tends to collapse with increasing osmotic pressure, while the semiflexible region remains extended. The centered pattern promotes hairpin-like conformations due to depletion interactions between the two semiflexible domains. Similar structures have been observed experimentally with actin filaments in crowded conditions, with depletion interactions promoting sustained contact between different ends of the same semiflexible polymer, resulting in a “racket-like” conformation. ^17^ The uniform pattern of flexibility results in more pronounced collapse of the polymer by facilitating interactions between multiple semiflexible domains. In general, large flexible regions tend to collapse with increasing crowding. However, when flexible regions separate large semiflexible domains, they can promote global collapse by facilitating depletion interactions between the semiflexible regions.

Figure 5 reveals key features of the adsorption behavior. At a given value of *η*, the diblock pattern adsorbs at the lowest levels of crowding, followed by the centered pattern. The uniform and random patterns of flexibility require higher osmotic pressures to adsorb. These trends can be understood in terms of the entropic costs of adsorption. Because there is a smaller loss of entropy upon adsorption of a semiflexible domain, long continuous semiflexible domains promote adsorption of the polymer to the surface. The longest semiflexible domains are found in the diblock pattern, followed by the centered pattern. Consistent with this result, continuous flexible domains maintain less contact with the surface than semiflexible domains at intermediate levels of crowding (Figure 6). These findings indicate that patterns of flexibility can be exploited to control polymer-surface interactions in crowded environments.

The osmotic pressures used in the simulations reside within experimentally accessible ranges achievable with common crowding agents such as poly(ethylene glycol) (PEG) or dextran. With PEG-8000 as a representative crowding molecule, we estimated crowder concentrations using scaled particle theory, ^56^ first calculating the volume fraction from the osmotic pressure, and then converting to a concentration using the specific volume. ^59,60^ For Π_*c*_ = 0.4, where strong adsorption was observed for some polymers, the corresponding volume fraction and concentration were 0.12 and 150 mg/mL, respectively. The highest crowding level studied, Π_*c*_ = 0.9, corresponded to a volume fraction of 0.20 and a concentration of 240 mg/mL.

The different collapse and adsorption behaviors observed in this study raise the question of whether spatially varying stiffness may fine-tune the conformations and organization of biological macromolecules, such as proteins and nucleic acids, in crowded cellular environments. In the context of cell-free biology, previous work has shown that crowding leads to surface association of DNA plasmids, which in turn affects transcription and translation.^14^ It would be interesting to explore whether sequence-dependent stiffness of DNA could be exploited to differentially control the surface association of different plasmids, thus enabling spatial control of transcription. Moreover, many soft-matter applications rely on controlled polymer conformations in crowded environments. DNA origami constitute an interesting class of nanomaterials in which relatively flexible regions can be used to connect stiff regions. Such DNA-based particles would provide a means to test how distributed flexibility impacts collapse and adsorption of particles, and the patterns of flexibility could be used to control adsorption behavior.

Overall, our results demonstrate that strategically arranging regions of different flexibility makes it possible to modulate polymer collapse and adsorption. Further understanding may enable the design of polymers with tailored collapse and adhesion characteristics, thus enabling new ways to control material properties.

## Data availability

The code used for this work, including custom LAMMPS files, simulation scripts, and analysis code, is available at https://github.com/GCantral/Crowding.

## Acknowledgements

This work was supported by the National Science Foundation (CBET-2217777). G. Cantrall was supported in part by a training grant from the National Institutes of Health (T32GM142621).

